# Computationally-guided design and affinity improvement of a protein binder targeting a specific site on HER2

**DOI:** 10.1101/2020.11.09.375618

**Authors:** Tae Yoon Kim, Jeong Seok Cha, Hoyoung Kim, Yoonjoo Choi, Hyun-Soo Cho, Hak-Sung Kim

## Abstract

A protein binder with a desired epitope and binding affinity is critical to the development of therapeutic agents. Here we present computationally-guided design and affinity improvement of a protein binder recognizing a specific site on domain IV of human epidermal growth factor receptor 2 (HER2). As a model, a protein scaffold composed of Leucine-rich repeat (LRR) modules was used. We designed protein binders which appear to bind a target site on domain IV using a computational method. Top 10 designs were expressed and tested with binding assays, and a lead with a low micro-molar binding affinity was selected. Binding affinity of the selected lead was further increased by two-orders of magnitude through mutual feedback between computational and experimental methods. The utility and potential of our approach was demonstrated by determining the binding interface of the developed protein binder through its crystal structure in complex with the HER2 domain IV.

## Introduction

A protein binder with a desirable epitope and a binding affinity is crucial to its development as a therapeutic agent since its efficacy is largely affected by a region where it binds on a target and its binding affinity (Ledford 2008; Chames et al. 2009). Experimental approaches comprising repeated rounds of a library construction and screening have been most widely used, but they are labor-intensive and time-consuming, and almost impossible to specify the binding region (Lerner 2006; Dunn 2010). The difficulty dramatically emerges if a target is a multi-domain protein with a large size. Especially, in the case of a library-based approach, selection of an initial binder usually determines the fate of a whole process including the *in vitro* and *in vivo* experiments.

Computational methods have recently attracted a considerable attention as a promising paradigm to design a protein binder with desired activity. Advances in computing power and algorithms have enabled the prediction of precise energy landscapes, leading to notable successes in computational protein designs (Silva et al. 2019; Chevalier et al. 2017; Ramisch et al. 2014; Tinberg et al. 2013; Fleishman, Whitehead, et al. 2011; Cannon et al. 2019). Despite many advances, however, purely computational design of a protein binder with a desired epitope and binding affinity remains a challenge. It has been known that current scoring functions may not be precise enough mainly due to limitations to accurately define the binding free energy landscapes (Houk and Liu 2017). Furthermore, if a target protein is composed of multi-domains and structurally flexible loops, it is extremely difficult to computationally design a protein binder with a desired epitope and a high affinity. Overall, design of such protein binder by computational method has been limited so far to target proteins with certain “ideal” features such as high secondary-structure content (Whitehead, Baker, and Fleishman 2013).

Human epidermal growth factor receptor 2 (HER2) is a well-known drug target for various cancers, representing a typical multi-domain membrane protein mainly composed of a number of flexible loops (Menard et al. 2003; Tebbutt, Pedersen, and Johns 2013; Cho et al. 2003; Banappagari, Ronald, and Satyanarayanajois 2010; Kastner et al. 2009). Monoclonal antibody trastuzumab is known to bind to the HER2 extracellular domain IV (HER2 domain IV), effectively inhibiting a HER2-mediated cell signaling process (Tebbutt, Pedersen, and Johns 2013; Arkhipov et al. 2013). Here we present computationally-guided design and affinity improvement of a protein binder targeting the trastuzumab epitope on domain IV of HER2 which mainly consists of flexible loops. As a model, a protein scaffold composed of LRR (Leucine-rich repeat) modules was employed. We firstly designed protein binders which appear to recognize the target site on domain IV through computational method based on the predicted complex model structures for HER2 domain IV and a protein scaffold (**Figure. 1**). Top 10 designs were expressed, and a lead with a low micromolar binding affinity was selected based on binding and inhibition assays. Binding affinity of the selected lead was further increased by two-orders of magnitude through mutual feedback between computational and experimental approaches. We demonstrated the utility of our approach by determining the binding interface of the developed protein binder through its crystal structure in complex with the HER2 domain IV. Details are reported herein.

**Figure 1.**
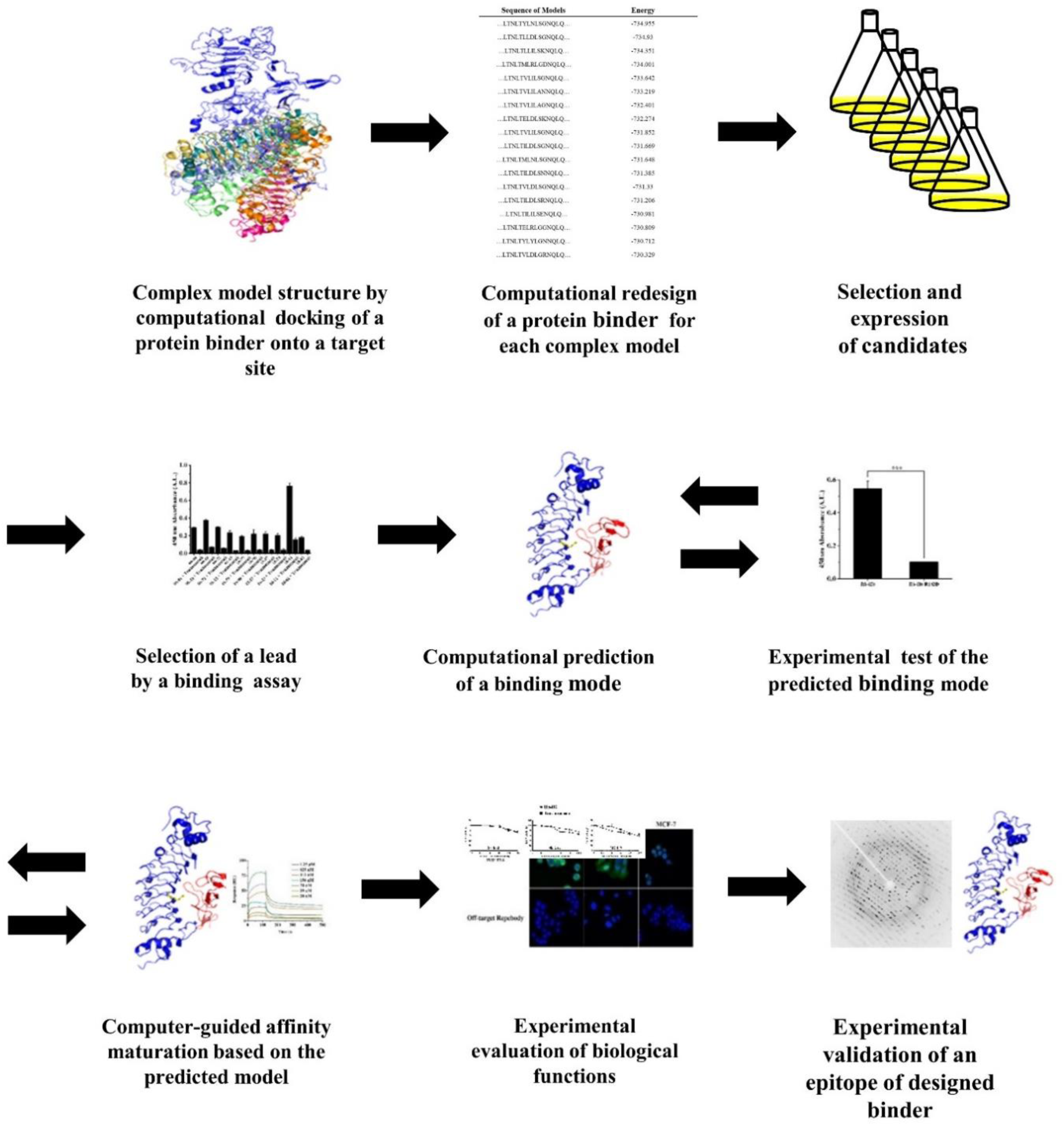
Schematic flow chart illustrating the computationally-guided design and affinity maturation of a protein binder targeting a specific site on HER2 extracellular domain IV.

## Results

### Computationally-guided design of protein binders targeting a specific site on HER2 domain IV

As proof-of-concept, we aimed to develop a protein binder which targets the trastuzumab epitope on HER2 domain IV and consequently inhibits a HER2-meidiated cell signaling. HER2 has no receptor ligands, and triggers cell signaling through homo- or heterodimerization with ErbB protein family (Baselga and Swain 2009). Interestingly, domain IV of all epidermal growth factor receptors is known to consist of structurally flexible loops (Banappagari, Ronald, and Satyanarayanajois 2010; Arkhipov et al. 2013), which may hinder computational design of a protein binder targeting such site. Monoclonal antibody trastuzumab was revealed to bind to the domain IV of HER2, and effectively inhibit a related cell-signaling process (Tebbutt, Pedersen, and Johns 2013; Arkhipov et al. 2013). Thus, while extremely challenging, potential therapeutic protein inhibitors should bind to a designated site of HER2 to have expected outcomes, when considering the molecular mechanism of the cell signaling. As a model, a protein scaffold composed of LRR (Leucine-rich repeat) modules, termed ‘Repebody’, was employed. The repebody scaffold showed desirable biochemical and physical properties such as high stability, easy module-based engineering, high bacterial expression, and high tissue penetration (Lee et al. 2012). A number of target-specific repebodies have been developed through phage display selection (Lee et al. 2014; Lee et al. 2015; Hwang et al. 2016). In particular, the target-binding region of the scaffold is composed of β-strands, exhibiting a rigid-body structure (Lee et al. 2012).

To successfully design a protein binder recognizing a target site through a computational method, the shape complementarity between a protein binder and a target site should be taken into consideration. As the shape complementary of a repebody may not exactly fit the entire trastuzumab epitope on HER2 domain IV, we estimated the chance of the overlap with the trastuzumab epitope by a repebody. Assuming that a library approach generates variants that can bind to any sites of HER2 domain IV, we generated 100,000 random docking models by assigning attraction on two nearby LRR modules, LRRV3 and LRRV4, of the repebody (**Figure 3a**). For this, the Rosetta docking protocol was employed (Weitzner et al. 2017). The simulation results show that 100 % overlap with the trastuzumab epitope using a repebody may not be possible (**Supplementary Figure S1**). In fact, nearly a half of the repebody models are not in contact with the epitope at all. Considering the shape complementarity and steric clashes against trastuzumab for inhibiting the cell signaling, an effective repebody should contain at least > 20 % of the trastuzumab epitope. We thus first generated complex model structures using ClusPro (Kozakov et al. 2017) with the antibody mode (Brenke et al. 2012) (while not particularly considering the trastuzumab epitope at this stage). Wild-type repebody (PDB ID: 3RFS) was docked onto the target site on domain IV of HER2 (PDB ID: 1N8Z) with repulsion constraints at HER2 domain I-III and the convex region of the repebody (**Figure 2a, b**). Total 30 docking models were generated for the target site on the HER2 domain IV. Each LRR module has four variable sites, and 20 variable sites in total on five modules of wild-type repebody were subjected to redesign based on the docking models using RosettaScript protocol (Fleishman, Leaver-Fay, et al. 2011). Only domain IV was considered in the Rosetta redesign process, and one thousand designs were generated for each docking model. Both proline and cysteine were excluded in the design process. Among the 30,000 designs, top 10 clones with the lowest energy values and those which appeared to share > 20 % of the trastuzumab epitope were selected. It should be noted that one of the initial docking models that share the epitope is very similar to the crystal structure of Rb-H2 in complex with the domain IV (**Supplementary Figure S2**). The selected clones were expressed in *E. coli* and subjected to purification, followed by binding assays using ELISA against the HER2 ectodomain (**Supplementary Figure S3**). We finally selected Rb-H0 as a lead showing the highest binding signal to HER2 ectodomain. There was a 5-fold signal decrease in the presence of trastuzumab (**Figure 2c**), which indicates that Rb-H0 shares the trastuzumab epitope. To estimate the binding affinity of Rb-H0, we analyzed the binding profile against HER2 using direct ELISA with the increasing concentration of Rb-H0. As a result, the binding affinity of Rb-H0 was estimated to be 4 μM (**Figure 3b**).

**Figure 2.**
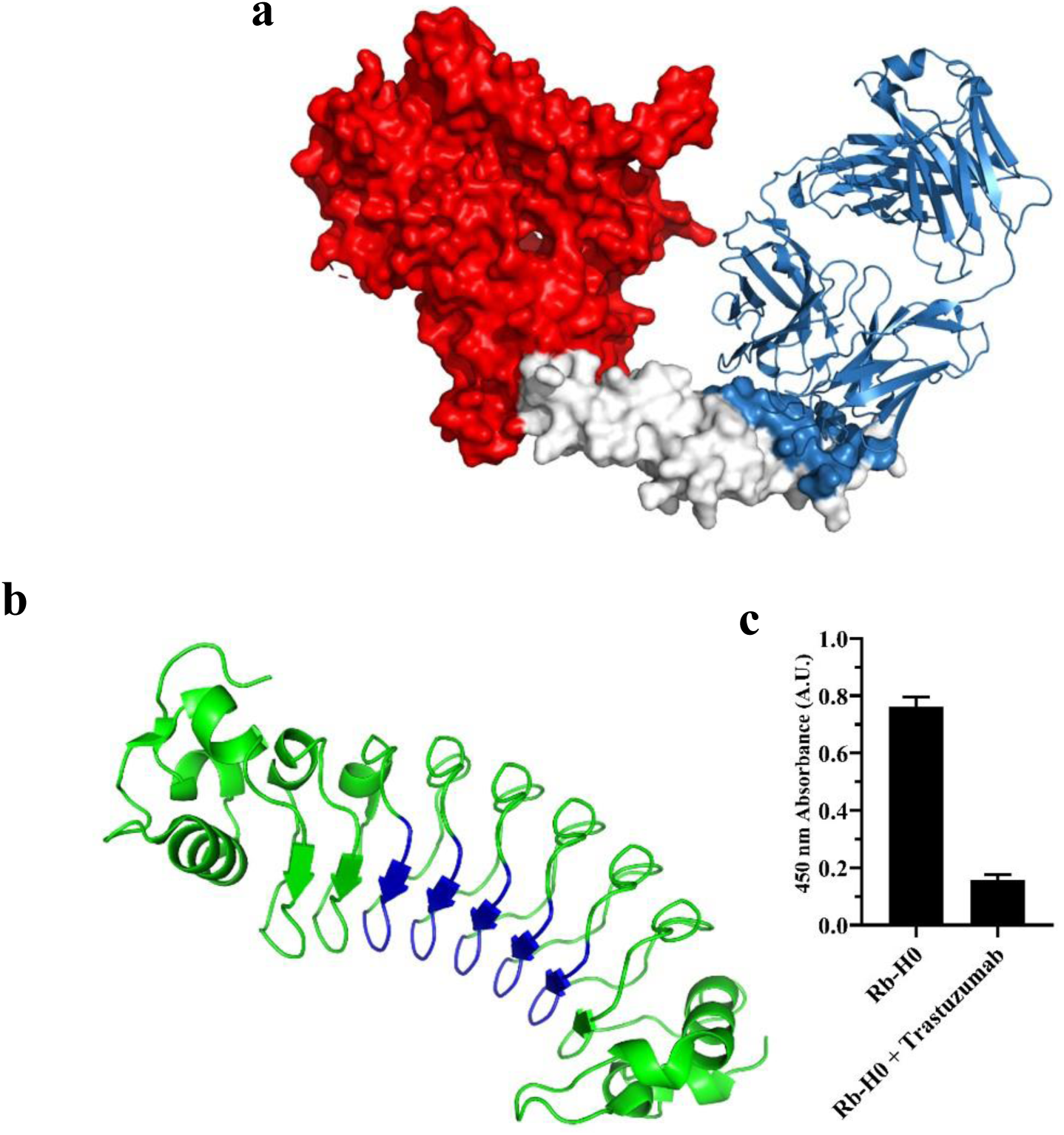
Selection of Rb-H0 binding to HER2 domain IV from the computationally designed protein binders. **a**, Initial docking models were generated on HER2 domain IV with assigned repulsion on domain I-III (red). Trastuzumab and its epitope are colored in skyblue. **b**, Non-concave region of a repebody model (PDB ID: 3RFS chain A) was masked (blue). **c**, Selection of Rb-H0 from the computationally designed candidates. Among top 10 designs showing the lowest energy levels, Rb-H0 exhibiting the highest signal and significant decrease in the signal in the presence of trastuzumab was selected as the initial binder. Error bars represent average ± standard deviation (n = 3).

**Figure 3.**
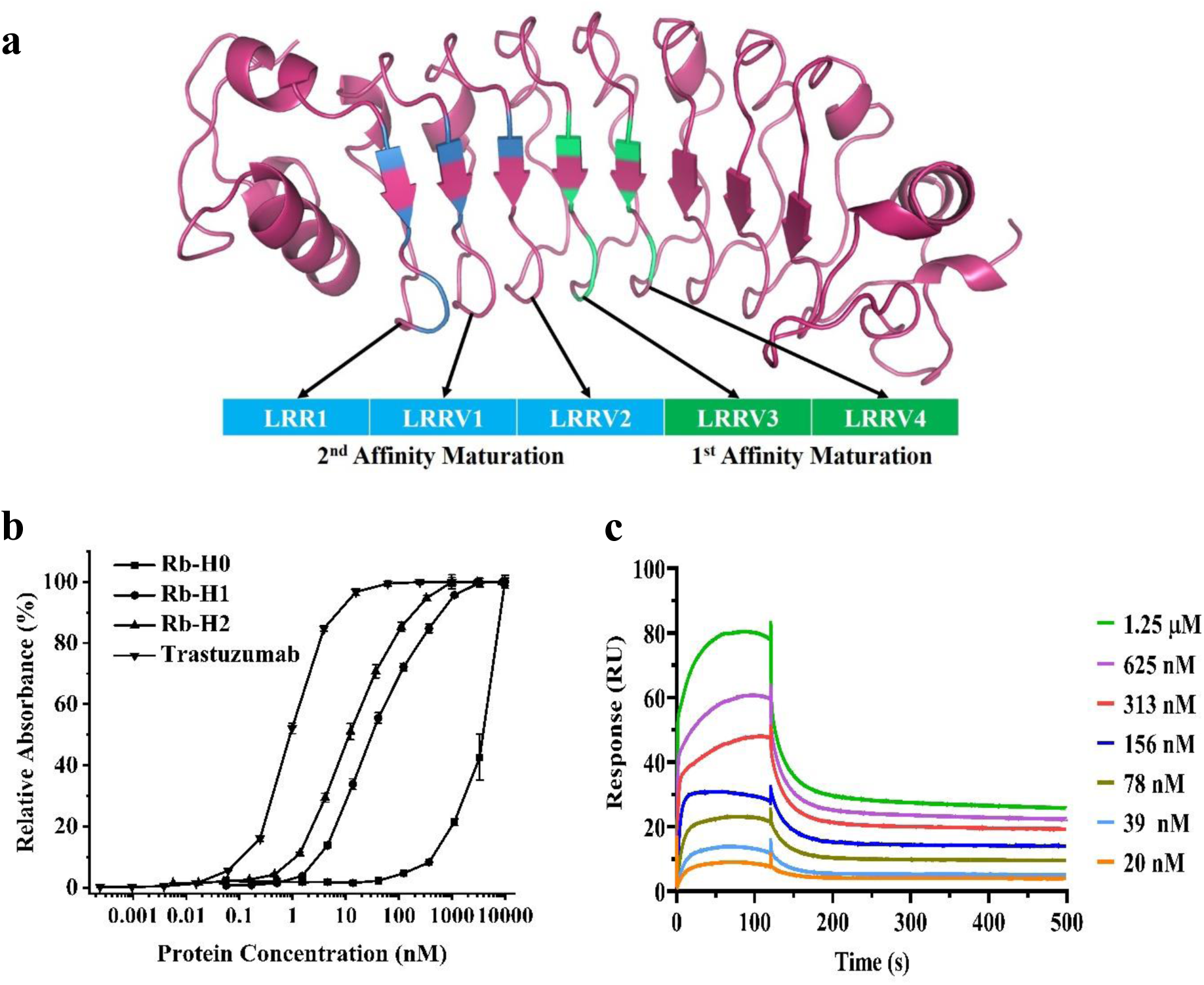
Computationally-guided affinity improvement of Rb-H0 and biophysical properties of affinity-maturated Rb-H2. **a**, Modules used in the first and second round of affinity maturation of Rb-H0 are shown in representative structure of a repebody scaffold. Annotation of the modules is indicated, and each module is numbered from N-terminus to C-terminus. Seven residues in modules LRRV3 and LRRV4 were used for first-round affinity maturation. Additional seven residues on modules LRR1, LRRV1, and LRRV2 were optimized for second round affinity maturation. **b**, Binding profiles of Rb-H0 and affinity-maturated Rb-H1 and Rb-H2 by ELISA. Error bars represent average ± standard deviation (n = 3). **c**, Determination of binding affinity of Rb-H2 for HER2 ectodomain through surface plasmon resonance (SPR).

### Affinity improvement by an integrated computational and experimental approach

Since Rb-H0 has a low binding affinity for HER2 ectodomain, we intended to increase its binding affinity. Based on the docking models of wild-type repebody against HER2 domain IV, we reasoned that the residues at modules LRRV3 and LRRV4 on Rb-H0 would have the highest proximity for the targeted site on HER2 domain IV. Seven residues (Ile114, Asp116, Ser118, Asn119, Ile138, Asp140, Ser142) were selected and randomized for a library construction followed by phage display selection (**Figure 3a**). A clone with the highest binding signal, designated as Rb-H1, was shown to have a significantly increased binding affinity compared with Rb-H0 (**Figure 3b**). For the second round of affinity maturation, we predicted the binding mode of Rb-H1 to the HER2 domain IV using the computational method as described elsewhere (Choi et al. 2019). It should be noted that the computational binding mode prediction requires solid experimental validation in advance (paratope information and clues on epitopes) and thus Rb-H0 binding mode prediction may not be performed. In the docking process, ClusPro with the antibody mode was employed for protein docking (Kozakov et al. 2017; Brenke et al. 2012). Rb-H1 structure was modeled based on wild-type repebody structure (PDB ID: 3RFS), and repulsion was assigned on the convex residues. Attraction was imposed on the library sites for affinity improvement. As it was known from the competitive binding assay that Rb-H1 might share the epitope with trastuzumab, any docking models that were not in contact with the trastuzumab epitope were eliminated. Total 17 docking models were finally selected, followed by energy-minimization using the Tinker molecular dynamics package (Rackers et al. 2018) (AMBER99sb (Hornak et al. 2006) with the GB/SA implicit solvent model (Still et al. 1990)), and the docking model with the lowest energy was predicted to be the binding mode of Rb-H1 (Choi et al. 2019) (**Supplementary Figure S4**). Based on the predicted binding mode, another seven residues (Gln46, Ile48, Asn50, Asn51, Tyr68, Ala70, Val90) on the three modules LRR1, LRRV1 and LRRV2, at the N-terminus were chosen and randomized for a library construction and phage display selection (**Figure 3a**). As a result, a variant with the highest signal, Rb-H2, was selected, and it was observed to have a marginal increase in binding affinity compared with Rb-H1 (**Figure 3b**). The binding affinity of Rb-H2 was determined to be 54 nM through surface plasmon resonance (SPR) (**Figure 3c**) which is a significant increase in the binding affinity of a lead protein binder by two-orders of magnitude.

After the second-round affinity-maturation, however, only a marginal increase in the binding affinity of Rb-H2 was observed despite seven additional mutations. Amino acid sequence of Rb-H0, Rb-H1, and Rb-H2 are shown in **Supplementary Table S1**. Given that information, we assumed that only certain mutations would contribute to the increase in the binding affinity of Rb-H1. We analyzed the binding energy for each single mutant of Rb-H2 based on the model complex for Rb-H1 (**Supplementary Table S2**). The energy calculation results indicate that two single mutations (V90T and N51H) make significant contributions to the increase in the binding affinity. By taking into account the calculation results, we remodeled the binding mode of Rb-H2 again by assigning attractions at the two predicted positions (V90T and N51H). After energy minimization, the final model was shown to be well coincident with the X-ray crystal structure (I-RMSD: 1.701 Å, f_nat_: 0.508, **Supplementary Figure S4**).

To further get insight into the binding site and affinity of the variants, competitive ELISA was carried out for Rb-H0, Rb-H1, and Rb-H2 in the presence of trastuzumab (**Figure 4a**). The signals of the variants were shown to decrease in the presence of trastuzumab, indicating that they share the trastuzumab epitope as intended and predicted. In the case of Rb-H2, the signal decrease was much smaller compared to Rb-H0, supporting a significant increase in binding affinity, considering the binding affinity of trastuzumab for HER2 domain IV (5 nM (‘Herceptin (Trastuzumab) [package insert]. U.S. Food and Drug Administration’ 1998)). We tested the specificity of Rb-H2 against ErbB family proteins with high structural similarity to HER2, including EGFR, HER2, HER3, and HER4. As a result, Rb-H2 showed the highest binding specificity for HER2 ectodomain (**Figure 4b**). Finally, we checked whether Rb-H2 forms a complex with HER2 ectodomain using size exclusion chromatography (**Figure 4c**). Rb-H2 in complex with HER2 ectodomain was eluted as a single peak at a designated position, confirming the complex formation between Rb-H2 and HER2 ectodomain. In addition, we double confirmed the complex formation between Rb-H2 and HER2 domain IV in gel filtration chromatography (Data not shown). The results support that Rb-H2 indeed shares a binding site on HER2 domain IV with trastuzumab as intended.

**Figure 4.**
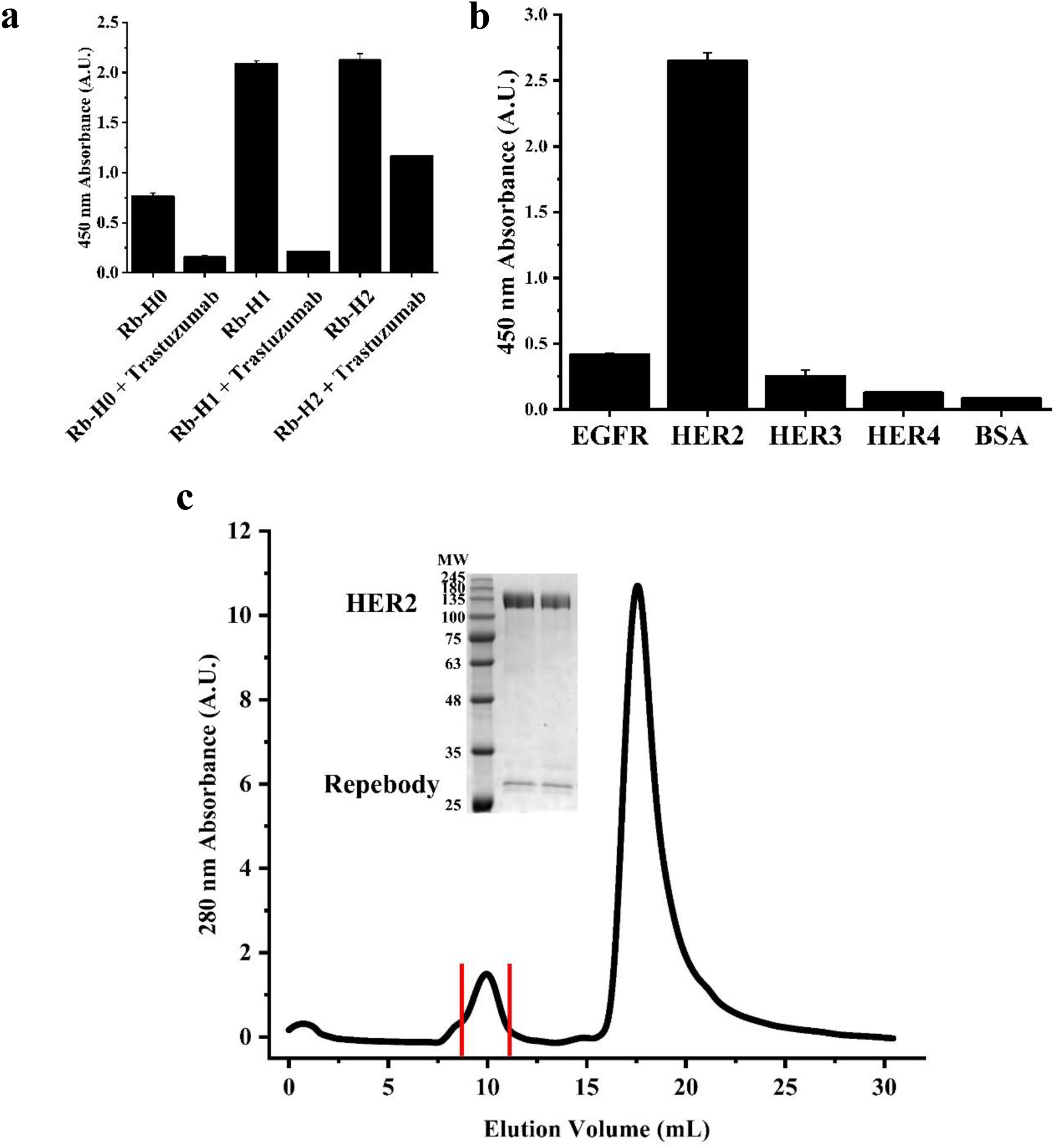
Characteristics of affinity-maturated Rb-H2. **a**, Competitive inhibition assays of Rb-H0, Rb-H1 and Rb-H2 in the presence of trastuzumab. All the binders showed decreased signals in the presence of trastuzumab. Error bars represent average ± standard deviation (n = 3). **b**, Specificity of Rb-H2 against ErbB family proteins. Rb-H2 was able to distinguish HER2 among the ErbB family proteins. Error bars represent average ± standard deviation (n = 3). **c**, Complex formation between Rb-H2 and HER2 ectodomain. Two proteins were mixed and eluted through size exclusion chromatography. Two proteins were eluted together in the first fraction as shown in SDS-PAGE (inset).

### X-ray crystal structure of Rb-H2 in complex with HER2 domain IV

To confirm the binding site of Rb-H2, we determined the X-ray crystal structure of Rb-H2 in complex with HER2 domain IV at 2.03 Å resolution (**Figure 5a, c**). The crystallographic and refinement statistics are shown in **Supplementary Table S3**. Rb-H2 is shown to bind to the targeted site of HER2 domain IV containing the trastuzumab epitope. The interface area between Rb-H2 and HER2 domain IV was estimated to be 2,070 Å^2^, whereas the interface area of trastuzumab is about 1,958 Å^2^. The binding site of Rb-H2 overlaps with that of the trastuzumab, covering approximately one-fourth of the trastuzumab epitope, which resulted in the binding competition against trastuzumab. Structural analysis revealed that HER2 domain IV interacts primarily with the concave side of Rb-H2 through hydrophobic interactions, hydrogen bonds and salt-bridge. Thr137 of Rb-H2 forms hydrogen bond with Gly572 of HER2 domain IV, and Trp116 and Arg140 of Rb-H2 have hydrophobic interactions with Val546, Leu547 and Val574 of HER2 domain IV (**Figure 5e**). In addition, Arg240 of Rb-H2 forms a salt bridge with Asp582 of HER2 domain IV, and four residues (Lys185, Tyr210, Arg240 and Gly244) of Rb-H2 have hydrogen bonds with four residues (Val585, Cys584, Asp582 and Gln583) of HER2 domain IV, respectively (**Figure 5f**). Specifically, three residues (Ser48, His51 and Tyr70) of Rb-H2 form hydrogen bonds with three residues (Glu543, Gln533, and Gln548) of HER2 domain IV (**Figure 5d**). Val162, Tyr210 and Trp212 of Rb-H2 are shown to have hydrophobic interactions with Ser573, Phe577 of HER2 domain IV (**Figure 5e**). It is interesting to note that some amino acid residues of Rb-H2 mentioned above (Ser48, His51, Tyr70, Trp116 and Arg140) are changed from those of Rb-H0. Overall, our structural analysis supports the significantly improved binding affinity of Rb-H2 for HER2 domain IV by two-orders of magnitude. Furthermore, based on the structure of HER2 domain IV, the binding region of Rb-H2 overlaps with the epitope of trastuzumab as shown in **Figure 5a** and **5b**. The binding surface areas of HER2 domain IV/Rb-H2, HER2 domain IV/ trastuzumab and the overlapped area are as follows: 2,070 Å^2^, 1,958 Å^2^ and 481 Å^2^ (23 % of the trastuzumab epitope), respectively. The overlapped residues between HER2 domain IV/Rb-H2 and HER2 domain IV/trastuzumab include Phe577, Asp582, Gln583 and Lys591 of HER2 (**Figure 5b)** (Cho et al. 2003). Based on our initial docking simulation (**Supplementary Figure S1**), the probability that a repebody generated from a random library approach shares > 23 % of the trastuzumab epitope is 0.3. The results provide distinct insight into the utility and potential of our computational-driven design approach.

**Figure 5.**
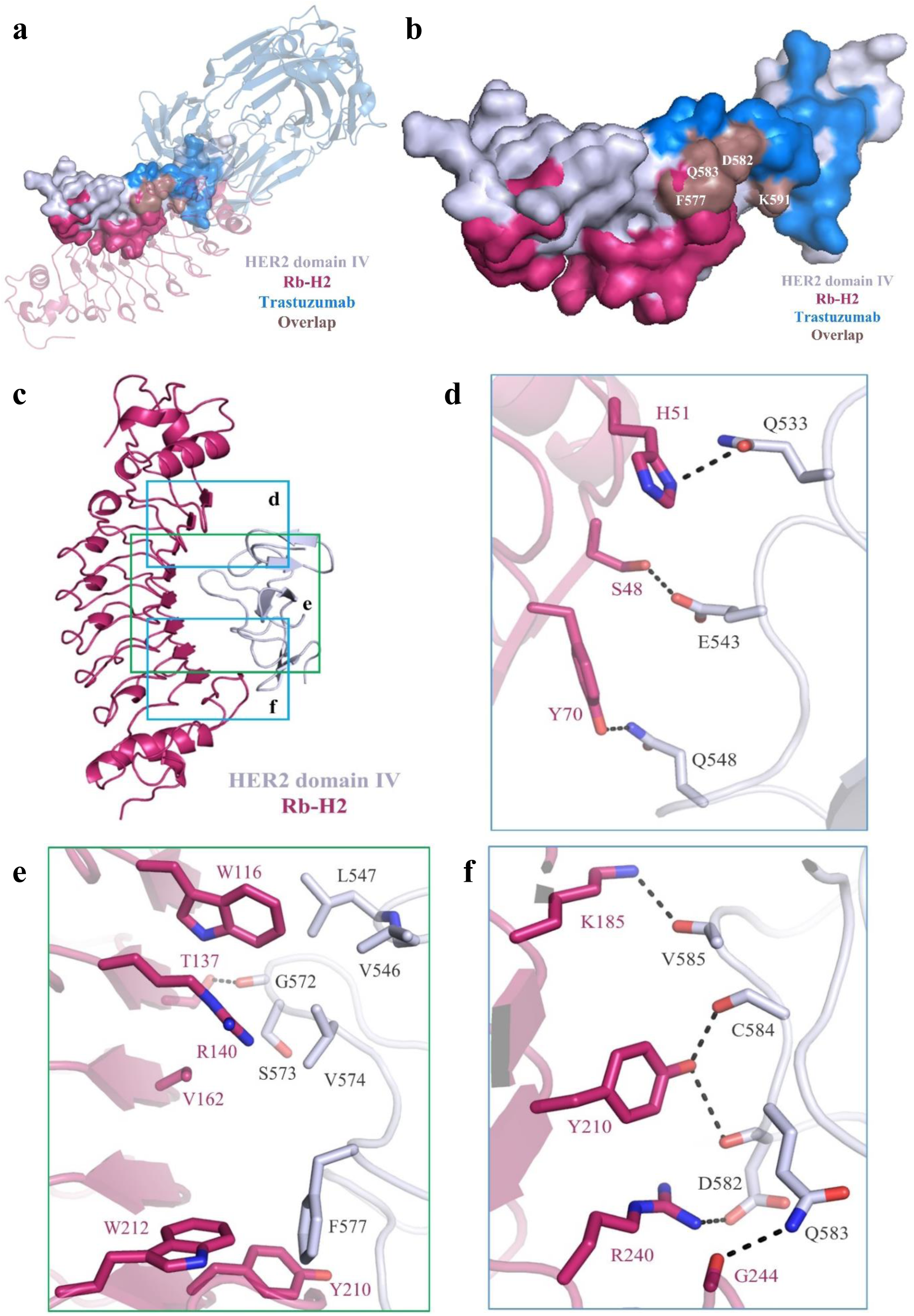
Crystal structure of Rb-H2 in complex with HER2 domain IV. **a**, Overall structure of Rb-H2 in complex with HER2 domain IV. The complex structure of trastuzumab in complex with HER2 domain IV is also shown. HER2 domain IV is presented in surface model, and Rb-H2 and trastuzumab are in cartoon model. Two structures are superimposed based on HER2 Domain IV. (HER2 domain IV: bluewhite, Rb-H2: warmpink, trastuzumab: skyblue, overlapped: dirtyviolet). **b**, Binding regions on HER2 domain IV of Rb-H2 and trastuzumab. The overlapped residues are marked. **c**, Overall structure of Rb-H2 in complex with HER2 domain IV. Rb-H2 and HER2 domain IV are presented in cartoon model. Three interaction regions are represented in detail in **d-f. d**, Hydrogen bonds are shown in dashed lines in stick model. **e**, Hydrogen bonds are shown in dashed lines in stick model and hydrophobic interaction residues are presented in stick model. **f**, Hydrogen bonds and salt-bridge are shown in dashed lines in stick model.

### *In vitro* binding and cytotoxicity of Rb-H2

We examined the binding of the developed Rb-H2 to HER2 on the cell surface. For this, cancer cell lines expressing different levels of HER2 were tested, including Sk-Br3 (high expression), Sk-Ov3 (moderate), and MCF-7 (low). We labeled Rb-H2 with fluorescein isothiocyanate (FITC) and treated it with the cells followed by imaging using confocal microscope. As shown in **Figure 6a**, the strong fluorescence intensity was observed on the peripheral region of Sk-Br3 cells, whereas MCF-7 cells exhibited the lowest fluorescent intensity. Based on the result, it is evident that Rb-H2 binds to HER2 ectodomain on the cell surface. No fluorescence was detected when an off-target repebody (human serum albumin specific repebody (Kim et al. 2019)) labeled with fluorescein was treated with each cell line, supporting the specific binding of Rb-H2 to HER2 ectodomain on the cell surface.

**Figure 6.**
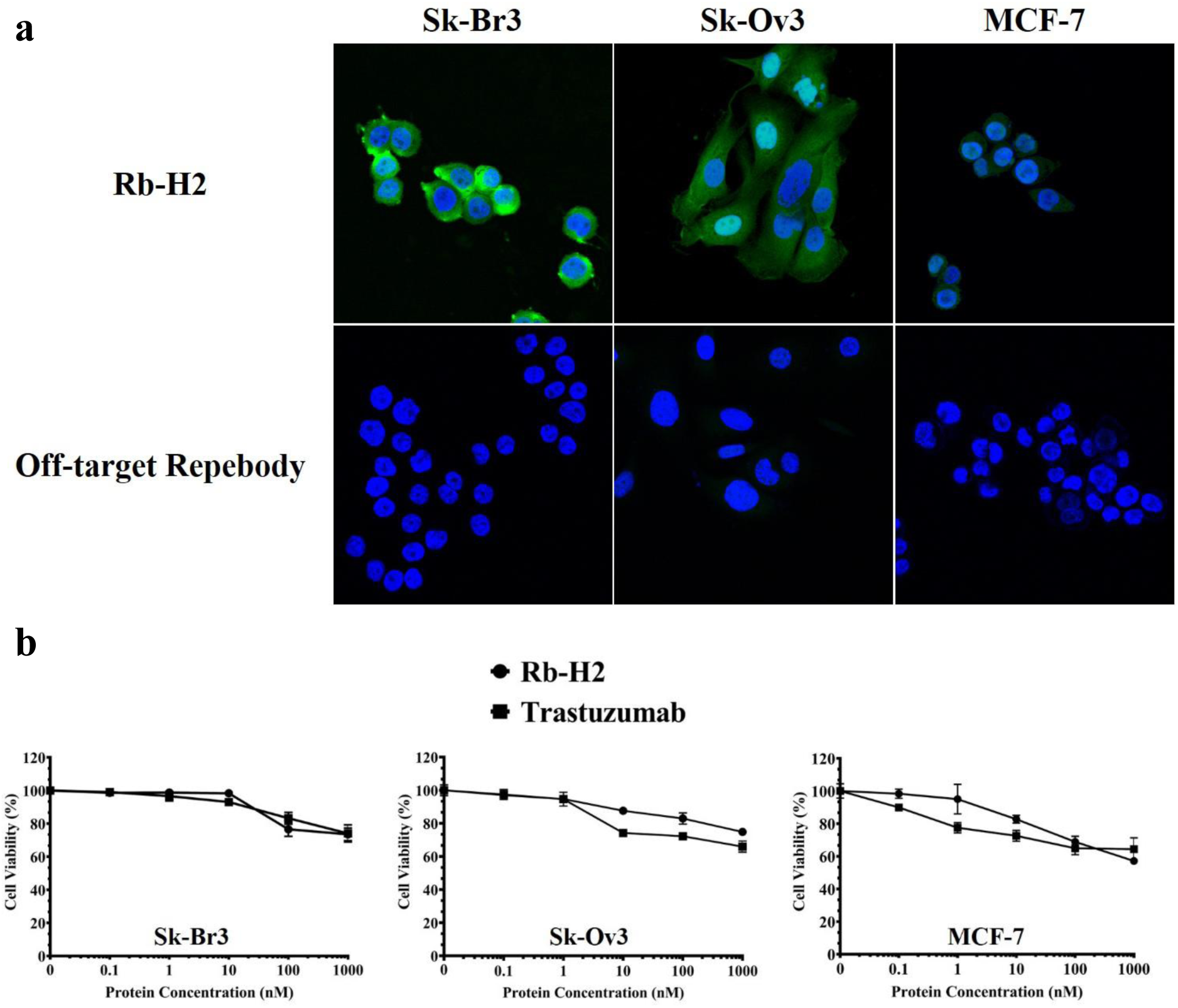
Binding of Rb-H2 to HER2-expressing cells and its cytotoxicity *in vitro*. **a**, Confocal microscopic images of HER2-expressing cancer cell lines after treatment with fluorescein-labeled Rb-H2. Cells were treated with 1 μM of labeled Rb-H2 for 3 h at 37 °C and imaged by confocal microscopy. Sk-Br3 (high level of HER2 expression, top), Sk-Ov3 (moderate HER2 level, middle), and MCF (low HER2 expression, down) cells were used. **b**, *In vitro* cytotoxicity of Rb-H2. Cells expressing HER2 were treated with Rb-H2 or trastuzumab at different concentrations for 72 h, and cell viability was determined by CCK-8 assay. Error bars represent average ± standard deviation (n = 3).

Since Rb-H2 was developed by targeting the trastuzumab epitope on HER2 domain IV, it is expected to inhibit the HER2-mediated cell signaling as trastuzumab does. We tested the cytotoxicity of Rb-H2 for various cancer cell lines (**Figure 6b**). In the case of Sk-Br3, cell viability gradually decreased with the increasing concentration of Rb-H2 and reached 70 % at the concentration of 1 μM. Similar cytotoxicity was observed for Sk-Ov3 and MCF-7, even though their HER2 expression levels were lower than Sk-Br3. Rb-H2 was shown to exhibit a similar cytotoxic pattern to trastuzumab for the tested cell lines, implying that it inhibits the cell signaling process in a similar way to trastuzumab because it shares the binding site with trastuzumab. Trastuzumab is clinically used for treatment of breast cancer and showed a saturation pattern even at high concentration. It is evident that Rb-H2 binds to the targeted site of HER2 domain IV and consequently inhibits the cell signaling pathway as trastuzumab by blocking dimerization, suppressing the cell proliferation.

## Discussion

We demonstrated computationally-guided design and affinity improvement of a protein binder recognizing a specific site on domain IV of HER2. Rational design of a protein binder with a desired epitope and binding affinity has been a long-standing goal in protein engineering field. Our strategy involves the computational design of protein binders which appeared to recognize a target site, followed by selection of potentially effective binders through experimental binding assays. As proof-of-concept, we aimed to design a protein binder which target the trastuzumab epitope on HER2 domain IV. The domain IV has very little content of secondary structures, which is supposed to be “non-ideal” features to be targeted by computational design approach. It has been shown that high flexibility of the domain makes it even harder to computationally design a protein binder recognizing such domain (Whitehead, Baker, and Fleishman 2013). Nonetheless, our approach enabled a successful design of a protein binder recognizing a target site on domain IV of HER2 as intended.

Epitope and binding affinity of a protein binder is crucial for its therapeutic efficacy. Development of a protein binder with a desirable epitope and binding affinity has mostly relied on experimental approaches comprising repeated rounds of a library construction and screening, but they are labor-intensive and difficult to identify the binding epitope during experiments. Recently, computational methods have shown notable successes in the design of proteins with desired functions due to many advances in the computing power and algorithms. However, purely computational design of a protein binder targeting a specific site still remains a challenge mainly because the current computational energy scoring is not accurate enough to precisely predict the binding free energy landscapes and may not be generally applicable. Furthermore, if a target protein is composed of multi-domains like extracellular receptors, computational design of such protein binders becomes extremely difficult. Our computationally-guided approach effectively generated the protein binder candidates for each docking model between a protein scaffold and HER2 domain IV by taking into consideration of shape complementarity. The docking models showed that a perfect overlap with the trastuzumab epitope using a repebody would be impossible as expected. Considering the shape complementarity and steric clashes against trastuzumab for inhibiting the cell signaling, we reasoned that an effective design should share at least > 20 % of the trastuzumab epitope. With this criterion, total 30 docking models were generated for the target site on the HER2 domain IV, and 20 variable sites of wild-type repebody were computationally redesigned for each model using RosettaScript protocol. Our computationally-guided approach eventually enabled a lead with a low micromolar affinity among the 30,000 designs. Based on the results, it is likely that shape complementarity is critical to the design of protein binder recognizing a target site. Experimental affinity improvement of a protein binder is generally known to be laborious and time-consuming. In contrast, our combined computational and experimental approach was shown to be effective for significantly enhancing the affinity of an initial binder, proving its utility for affinity improvement.

The X-ray crystal structure of Rb-H2 in complex with HER2 domain IV validated the utility of our approach by confirming that Rb-H2 indeed binds to the target site on HER2 domain IV as intended, showing the overlap (approximately one fourth) with the trastuzumab epitope. It is interesting to note that the computationally predicted binding orientation of Rb-H2 against HER2 domain IV was well coincident with the X-ray crystal structure, which supports the utility of the computational method to model the binding mode (Choi et al. 2019). Binding of Rb-H2 to the HER2-expressing cells and its *in vitro* cytotoxicity also supported the potential of our approach. Taken together, the present strategy can be widely applied to the development of a protein binder with a desired epitope and binding affinity for a target protein as an alternative to conventional experimental methods.

## Materials and Methods

### Synthesis and expression of genes

Computationally designed repebody genes and primers used for phage display library were synthesized from Integrated DNA Technologies (Coralville, IA, USA). Synthesized gene fragments went through cloning process after overnight digestion with restriction enzymes (Nde I, Xho I) at 37 °C and ligation (T4 DNA Ligase, Takara Bio, Shiga, Japan) into pET21 vector (Novagen, Madison, WI, USA) at room temperature for 2h. Materials for bacterial culture were supplied from Duchefa (Haarlem, The Netherlands). Origami B (DE3) competent cells (Novagen) were used for repebody expression. Isopropyl β-D-1-thiogalactopyranoside (IPTG) was purchased from LPS Solution (Seoul, Korea). The Ni-NTA agarose resin for purification of his-tagged proteins was purchased from Qiagen (Germantown, MD, USA). Superdex 75 16/600 and Superdex 200 Increase 10/300 size exclusion chromatography columns were purchased from GE Healthcare (Uppsala, Sweden). All other reagents including buffers and solvents were of analytical grade.

### Phage display selection

A repebody library was constructed by overlap PCR using primers containing NNK codon for variable sites on each module. The resulting library was inserted into pBEL118N phagemid (Lee et al. 2012) and electroporated to TG1 Electroporation-Competent Cells (Agilent Technologies, Santa Clara, CA, USA). Phages containing the library were rescued using M13KO7 Helper Phage (New England Biolabs, Ipswich, MA, USA). Solution-phase bio-panning was conducted in order to minimize the disruption of a target protein. Briefly, 10 μL of Dynabeads M-280 Streptavidin (Invitrogen, Waltham, MA, USA) was loaded into a sterile 1.5 mL centrifuge tube for immobilization of a target protein, and 40 μL of Dynabeads were added to another tube for a negative selection. After washing the beads with PBS (pH.7.4) twice by brief vortex, biotinylated HER2 ectodomain (4 μg/mL, Sino Biological, Beijing, China) was added to the tube and incubated for 2 h at 4 °C. Both HER2 ectodomain-bound beads and negative selection beads were blocked with PBST (PBS pH 7.4, 0.05% Tween 20) containing 2% BSA for 2 h at 4 °C. Phages were prepared in PBST containing in 1% BSA at a final phage concentration of 1.0 × 10^12^ cfu/mL and mixed with negative selection beads for 1 h. Purified Rb-H0 or Rb-H1 protein was also added to the phage solution in 1 µM concentration for competition. Phage solution was separated from the beads using a magnet and added to HER2 ectodomain-bound beads. After 2 h incubation at room temperature, beads were isolated by using magnetic bar and incubated for 1 min with PBST. This process was repeated five times and finally washed with PBS. To disrupt the binding between a target protein and phage-displayed repebody, 0.2 M glycine-HCl solution (pH 2.2) was added. Beads were isolated by a magnetic bar, and 1M Tris solution (pH 9.0) was added to the supernatant for neutralization before mixing with TG-1 cells. Phage-infected TG-1 cells were grown in 2xYT agar plate supplemented with 100 μg/mL ampicillin and 1% glucose for overnight at 30 °C. On the following day, 2xYT media were added into the plates to gather the cells and used for next round of selection process. Total five rounds of selection process were conducted for enrichment of positive clones. After the 5^th^ round, cells were diluted at appropriated ratio with PBS (pH 7.4) before plated. On the following day, a 96 deep-well plate (Axygen Scientific, Corning, NY, USA) was seed with colonies and the resulting phages were acquired for phage ELISA as described in our previous work (Kim et al. 2019). Phages were detected by HRP-conjugated anti-M13 antibody (GE Healthcare). For signal generation, 3,3’ 5,5’-tetramethylbenzidine (TMB) (Sigma Aldrich, St. Louis, MO, USA) was used, and the reaction was stopped by addition of 1N H_2_SO_4_. Absorbance at 450 nm was measured with Infinite M200 microplate reader (Tecan, Crailsheim, Germany).

### Computational design and affinity maturation of a site-specific repebody

Initial binding orientations were generated using the ClusPro webserver with the antibody mode (Brenke et al. 2012). Wild-type repebody (PDB code: 3RFS) was used as a receptor, and the crystal structure of HER2 (1N8Z) was employed as a ligand. The amino acid residues at the convex region were assigned to be repulsive. The residues of the trastuzumab epitope on HER2 domain IV were defined using PyMol from the X-ray crystal structure of trastuzumab in complex with HER2 ectodomain (1N8Z). Any residues on HER2 ectodomain within 5 Å from trastuzumab were defined as epitopes, and attraction was imposed. As a result, 30 binding models were generated. Rosetta 3.6 (2016.34) with talaris2014 was employed to redesign the binding sites of a repebody toward HER2 domain IV for each binding model (Fleishman, Leaver-Fay, et al. 2011; O’Meara et al. 2015). All amino acid types except for cysteine and proline were allowed at each position. For each model, 1,000 designs were generated, and the top 10 designs with the lowest energy values among the 30,000 designs were selected for further tests. The binding mode prediction was performed using the TINKER molecular dynamics package (Rackers et al. 2018). The AMBER99sb with the GB/SA implicit solvent model was used to minimize the model (Hornak et al. 2006; Still et al. 1990). For computer-guided affinity maturation, the residues were selected to increase the interaction with HER2 domain IV based on predicted binding modes. Selected residues were randomized to generate a library for phage display, and a clone with highest binding affinity was selected using phage ELISA. The same procedure was repeated to further increase the binding affinity of a selected repebody.

### Enzyme-linked immune-sorbent assay (ELISA)

Binding property of designed and selected repebodies were analyzed by direct ELISA. Briefly, a 96-well Maxibinding plate was coated with extracellular domain of HER2 at 4 °C overnight. PBST containing 1% BSA was used for blocking and dilution of repebodies and antibodies. PBST was used as washing buffer throughout the process. The repebody was detected by using HRP-conjugated anti-c-Myc antibody (1:500 dilution, Santa Cruz Biotechnology, Dallas, TX, USA) or biotinylated anti-repebody antibody (1 μg/mL, AbClon, Seoul, Korea) and HRP-conjugated streptavidin (1:1000 dilution, BioLegend, San Diego, CA, USA). For trastuzumab (Herceptin), HRP-conjugated anti-human Fc antibody (1:10000 dilution, Sigma Aldrich) was used. TMB solution was used for a signal generation and the reaction was stopped using 1N H_2_SO_4._ The signals were measured at 450 nm by microplate reader, and absorbance from maximum concentration was converted to 100 % for comparison. For binding specificity test against ErbB family proteins, EGFR, HER3 and HER4 proteins were used (Sino Biological).

### Surface plasmon resonance (SPR)

Binding affinity of a repebody was determined through surface plasmon resonance (Biacore T200, GE Healthcare). Briefly, 250 μg/mL of NeutrAvidin Protein (Thermo Scientific, Waltham, USA) was first coated on the surface of CM5 chip (GE Healthcare) in 10 mM sodium acetate buffer (pH 4.5). After immobilization, 20 μg/mL of biotinylated HER2 ectodomain was injected into the chip. Sensograms were obtained by flowing a serially diluted repebody into the chip. Kinetic constants were determined by the 1:1 Langmuir binding model using Biacore T200 software (GE Healthcare).

### Size exclusion chromatography for the complex formation

20 μg of HER2 ectodomain from Abcam (Cambridge, UK) was mixed with 5-fold excess amount of a repebody and incubated at 4 °C overnight. The mixtures were injected into Superdex 200 increase 10/300 column for analysis. The peak fractions were analyzed by SDS-PAGE.

### Expression and purification of HER2 domain IV

HER2 domain IV which corresponds to residues from 531 to 626 of HER2 was expressed using insect cells. HER2 gene was subcloned into a baculovirus expression vector by adding Mellitin signal peptide sequences and nona-histidine tag to the N-terminal of HER2 domain IV and TEV cleavage site and maltose-binding protein (MBP) tag to the C-terminal of HER2 domain IV. The expression of HER2 domain IV in insect cells was carried out using a Bac-to-Bac® Baculovirus Expression System (Invitrogen). The resulting construct was expressed in *Spodoptera frugiperda* (Sf9) insect cells in a secreted form through a culture at 27°C for 3 days. The media containing secreted HER2 domain IV were collected through centrifugation to remove Sf9 cells and adjusted to a pH of 7.5 for filtration before purification. Next, HER2 domain IV was purified using a HisTrap excel column (GE Healthcare). The filtrated media were loaded into the HisTrap excel column, followed by washing with 20 mM Tris-HCl (pH 7.5), 100 mM NaCl and 20 mM Imidazole. Bound HER2 domain IV was eluted with an elution buffer (20 mM Tris-HCl, pH 7.5) containing 100 mM NaCl and 250 mM Imidazole. Thereafter, the TEV recognition site was cleaved using TEV protease. After desalting to 20 mM Tris-HCl (pH 7.5) containing 50 mM NaCl, HER2 domain IV was loaded into an anion-exchange chromatography column (HiTrap-Q, GE healthcare), and HER2 domain IV was collected from flow through.

### Crystallization, data collection, and structure determination of Rb-H2 in complex with HER2 domain IV

Rb-H2 and HER2 domain IV were mixed at a 1:1.5 molar ratio and incubated for 1 h at 4°C. The mixture was applied to the size-exclusion column (HiLoad 16/600 Superdex 200 pg, GE Healthcare). The complex protein between Rb-H2 and HER2 domain IV was concentrated at up to 10 mg/ml and used for crystallization. Initial crystallization screening was conducted by using Mosquito robot (TTP Labtech, Melbourn, UK), and single, appropriate size of crystals appeared at 0.1 M Sodium Citrate: Citric Acid (pH 5.5) and 20% PEG 3000. The complex crystals were quickly soaked into a crystal buffer containing 20 % ethylene glycol to protect the crystals from the low temperature of the liquid nitrogen. X-ray diffraction of the complex crystal was then conducted to collect diffraction images using a BL-1A micro-beam line at the Photon Factory (Japan). An integration of the images was conducted using the XDSGUI, and a scaling of the mtz file was also performed using the CCP4 program (Winn et al. 2011). The complex crystal belongs to the space group P212121 with a = 44.66 Å, b = 80.07 Å, and c = 108.41 Å in a cell unit. The initial phase was obtained through a molecular replacement (MR) by Molrep using HER2 (PDB ID: 1N8Z) and repebody (PDB ID: 5B4P) as the initial searching model(Vagin and Isupov 2001). Model building was conducted using the Coot program, and refinement was achieved using Refmac5 (Emsley et al. 2010; Murshudov, Vagin, and Dodson 1997). A three-dimensional representation of the structure was carried out using the PyMOL program.

### Cell culture

Sk-Br3, Sk-Ov3, MCF-7, MDA-MB-468 (ATCC, Manassas, VA, USA) cell lines were cultured in RPMI 1640 media supplemented with 10% FBS, 100 U/mL penicillin, 100 μg/mL streptomycin (Capricorn Scientific, Ebsdorfergrund, Germany) at 37 °C incubator with 5% CO_2_.

### Immunofluorescence labeling and confocal microscopy

NHS-Fluorescein (Thermo Scientific) was prepared in DMSO (Sigma Aldrich) at a concentration of 10 mg/mL and mixed with a repebody dissolved in PBS (pH 7.4) at a dye-to-protein ratio of 10 with a final concentration of a repebody aimed to 2 mg/mL. The mixture of protein and dye was incubated at 4 °C overnight. Excess dye was removed using 0.22 μm centrifugal filter at 13000 rpm for 10 min and subjected to PD-10 desalting column (GE Healthcare). The concentration of a FITC-labeled repebody was measured by NanoDrop 2000c (Thermo Scientific). For confocal microscopy, cells were detached using non-enzymatic cell dissociation solution (Sigma Aldrich) when they reached 80 % confluence, and seeded into a 8-well slide glass (SPL Life Sciences) at 3.5×10^3^ cells/well. After 48 h of incubation, cells were gently washed with DPBS (Welgene, Seoul, Korea) for 3 times and treated with a FITC-labeled repebody at 4 °C to prevent endocytosis for 2 h. Following the removal of proteins, cells were gently washed again with DPBS for 3 times and fixed with 4 % paraformaldehyde in PBS for 30 min at room temperature. After washing with DPBS for three times, cells were stained with DAPI. Cell images were obtained using LSM 780 Confocal Microscopy (Carl Zeiss, Oberkochen, Germany).

### *In vitro* cytotoxicity

Cells were detached using trypsin-EDTA (Gibco, Waltham, MA, USA) when they reached 80 % confluence and seeded into a 96-well plate (SPL Life Sciences) at 1×10^4^ cells/well. After 24 h of incubation, cells were treated with serially diluted (10-fold) repebody or trastuzumab in RPMI 1640 serum free media and further incubated for 72 h at 37 °C and 5 % CO_2_ chamber. Cell cytotoxicity was measured by Cell Counting Kit-8 (Dojindo Molecular Technologies, Kumamoto, Japan). Signals were detected at 450 nm using Infinite M200 microplate reader. Absorbance from cells treated with only media was converted to 100 % for comparison.

## Acknowledgements

This research was supported by the Bio & Medical Technology Development Program (NRF-2017M3A9F5031419 to H.-S.K., NRF-2017M3A9F6029755, NRF-2019M3E5D6063903 to H.-S.C), Global Research Laboratory (NRF-2015K1A1A2033346 to H.-S.K.), Mid-Career Researcher Program (NRF-2017R1A2A1A05001091 to H.-S.K.), NRF-2018R1A5A2024181 to Y.j.C., Science Research Center (NRF-2016R1A5A1010764 to H.-S.C) of the National Research Foundation (NRF) funded by the Ministry of Science and ICT of Korea. We thank the staff scientists for assistance at the beamline 1A and 17A of the Photon Factory and the beamline 11C of Pohang Light Source.

## Data availability

Protein structure information are deposited in Protein Data Bank (Accession code: 6LBX).

