## Supplementary Information for "Computationally-guided design and affinity improvement of a protein binder targeting a specific site on HER2"

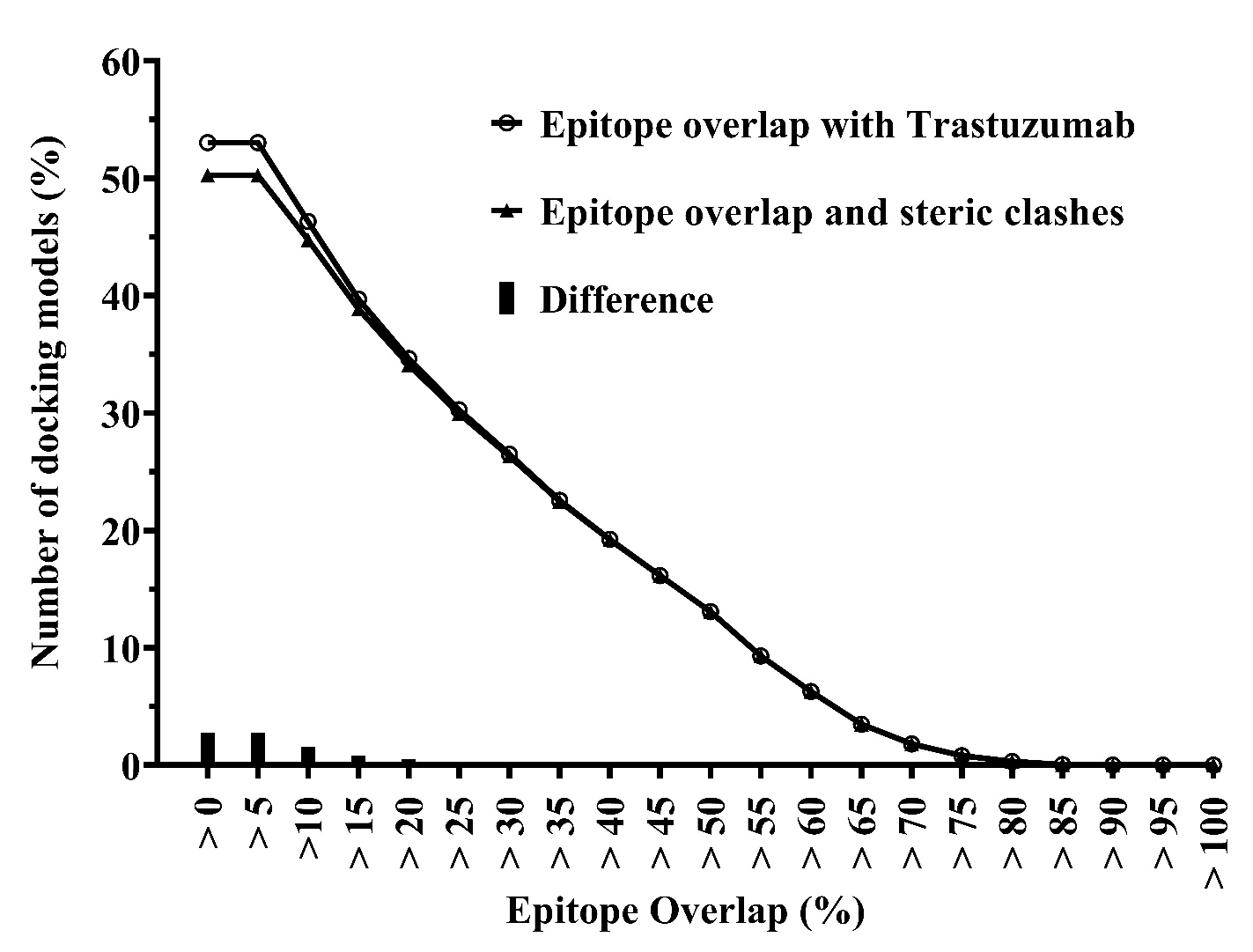

**Supplementary Figure S1.** Docking simulations for the repebody shape complementary. The Rosetta docking protocol generated 100,000 random docking models for wild-type repebodies on HER2 domain IV. Approximately a half of the models are not in contact with HER2 domain IV, and there is no model that cover the entire trastuzumab epitope. There exist some docking models that share the epitope but no steric clashes with trastuzumab (the black bar). If the epitope overlap is over 20 %, > 99.5 % of the docking models may physically disrupt the binding of trastuzumab to HER2 domain IV.

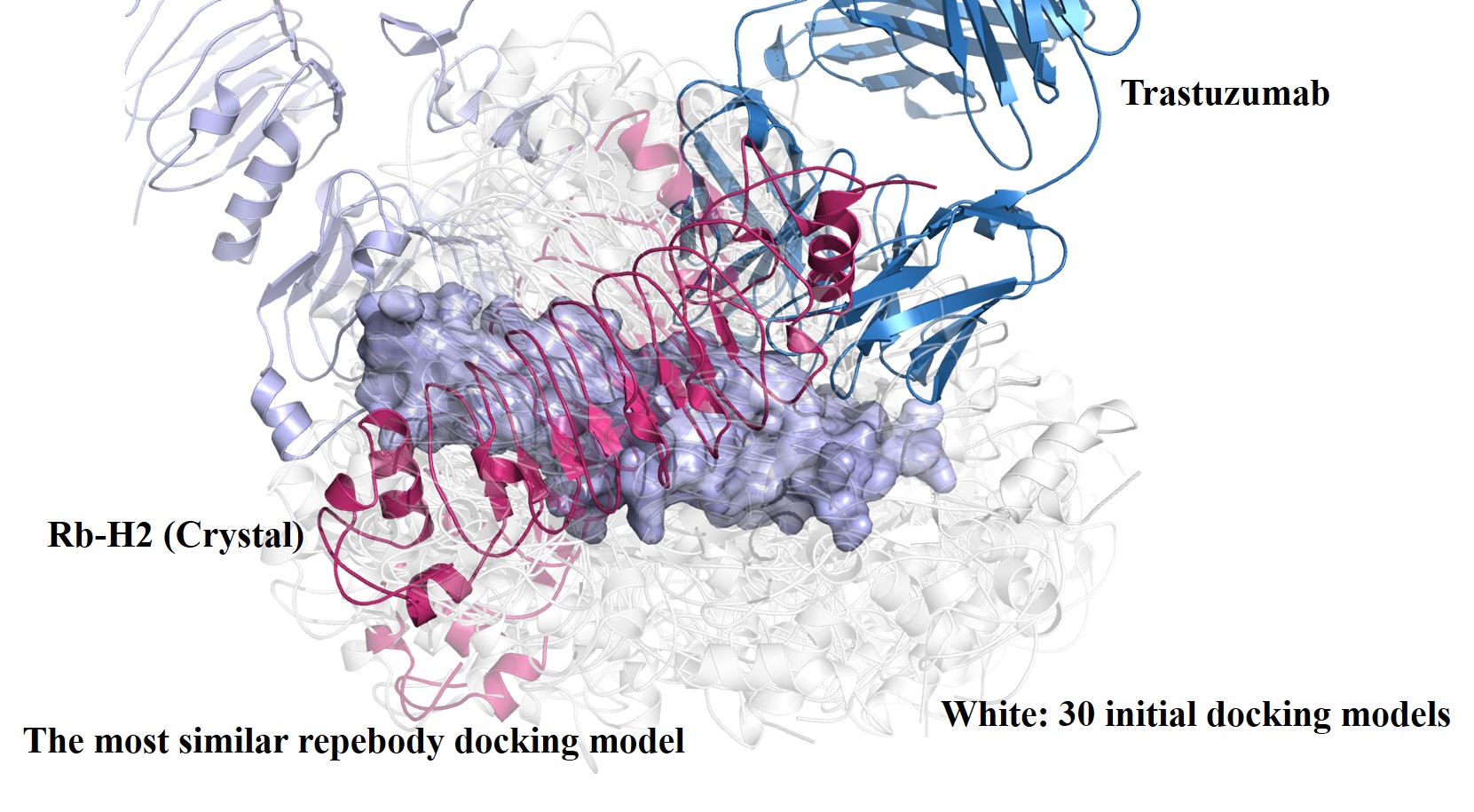

**Supplementary Figure S2.** Initial docking models. Among the 30 initial docking models, one of the initial docking models that share the epitope was similar to the crystal complex structure of Rb-H2 and the domain IV.

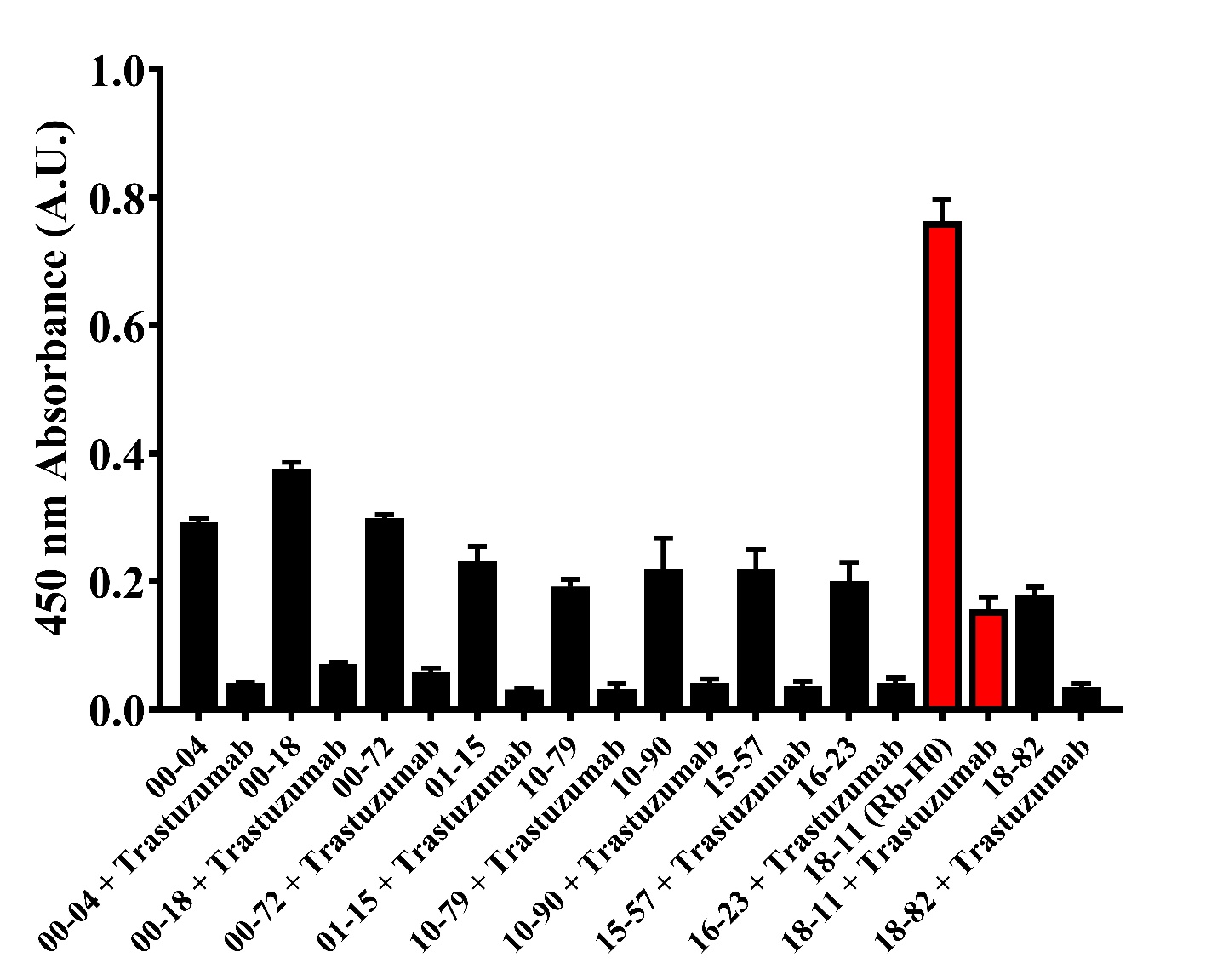

**Supplementary Figure S3.** Binding profile and trastuzumab competition level of top 10 designed candidates with the lowest energy was evaluated by direct ELISA. The repebody clone 18-11 showed highest signal and significant decrease of signal by trastuzumab was selected as the initial binder (Rb-H0) and subjected to further optimization.

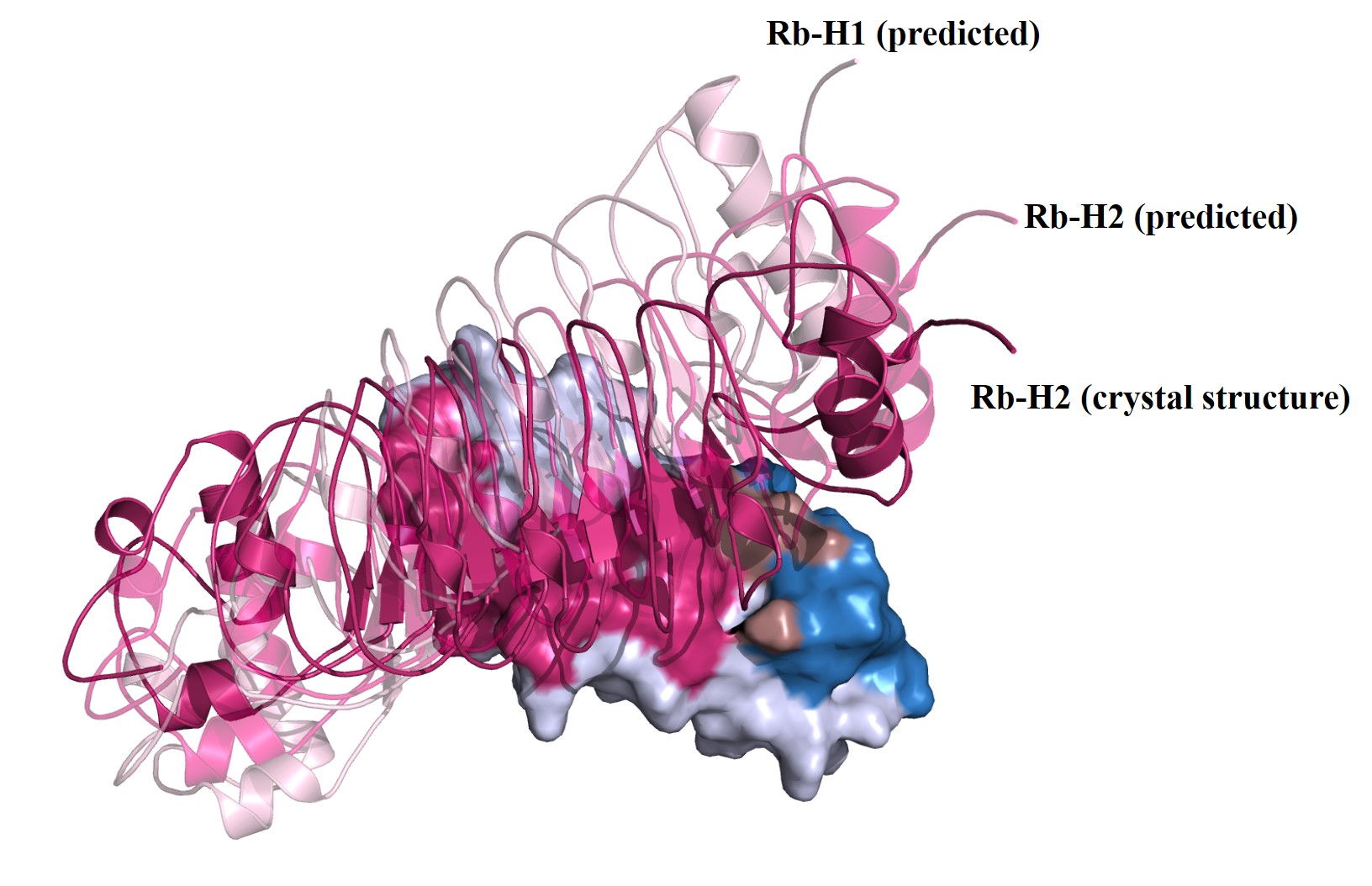

**Supplementary Figure S4.** Comparison of binding profile between Rb-H0 (predicted, white), Rb-H1(predicted, pale warmpink), Rb-H2 (predicted, light warmpink) and Rb-H2 (crystal, warmpink) against HER2 domain IV (bluewhite). The trastuzumab epitope and Rb-H2 are colored in skyblue and warmpink, respectively, on HER2 domain IV.

**Supplementary Table S1.** Complete amino acid **s**equences of Rb-H0, Rb-H1 and Rb-H2. Amino acid residues of Rb-H0 selected for library generation are colored in red.

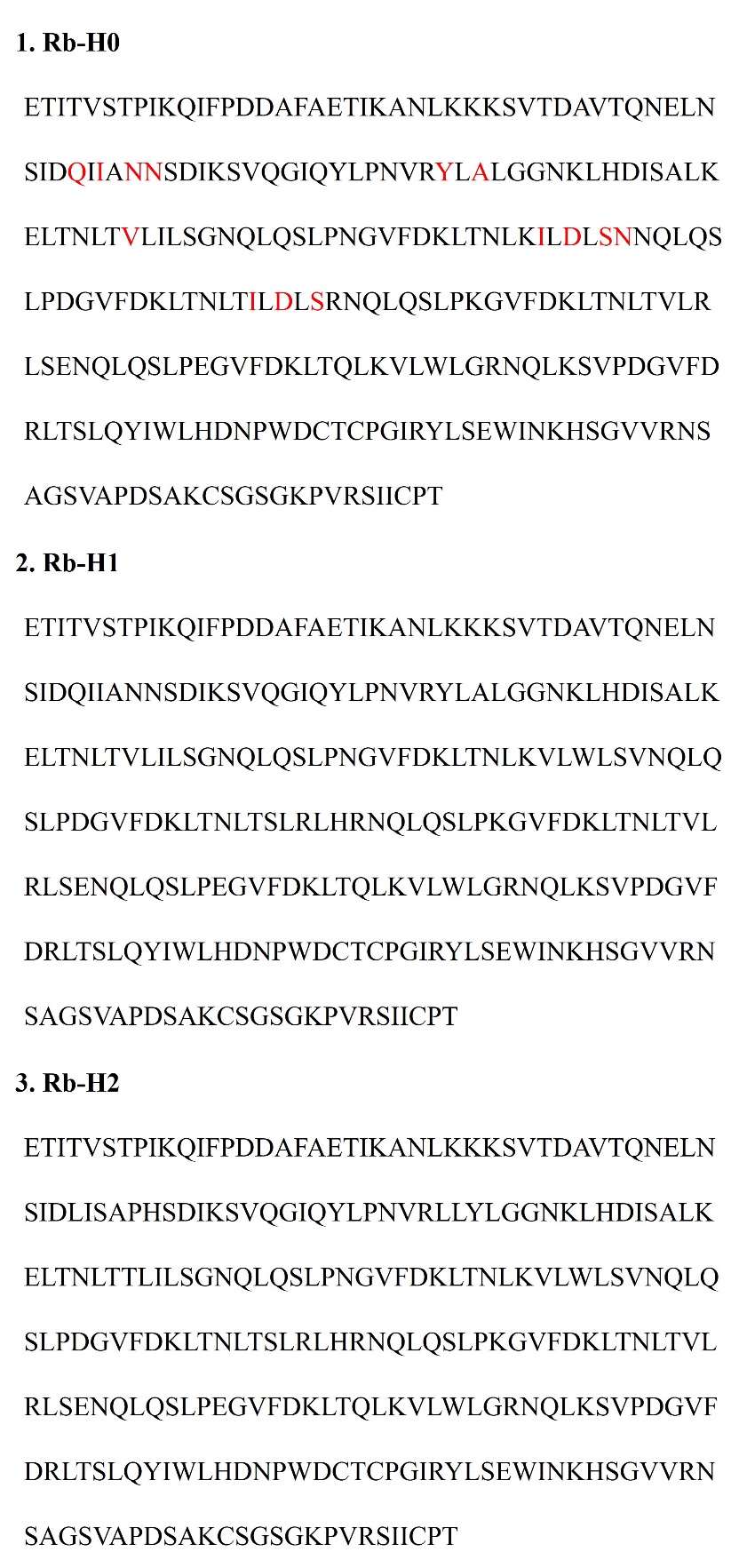

**Supplementary Table S2.** Differences in intermolecular energy values calculated for single mutation on Rb-H2 based on the model complex for Rb-H1. Two mutations, Val90Thr, Asn51His, were predicted to make significant contributions to an increase in binding affinity for HER2 domain IV.

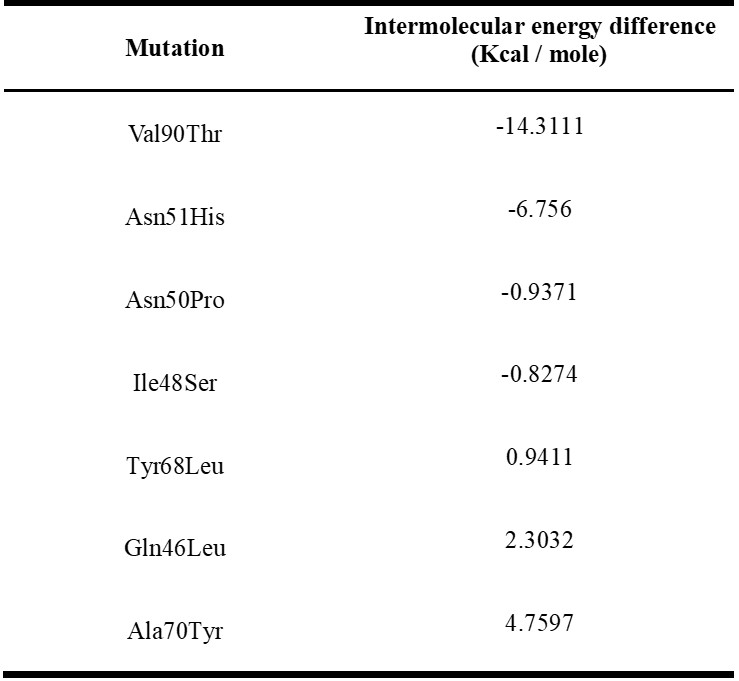

**Supplementary Table S3.** Parameters for crystal structure of HER2 domain IV and Rb-H2.

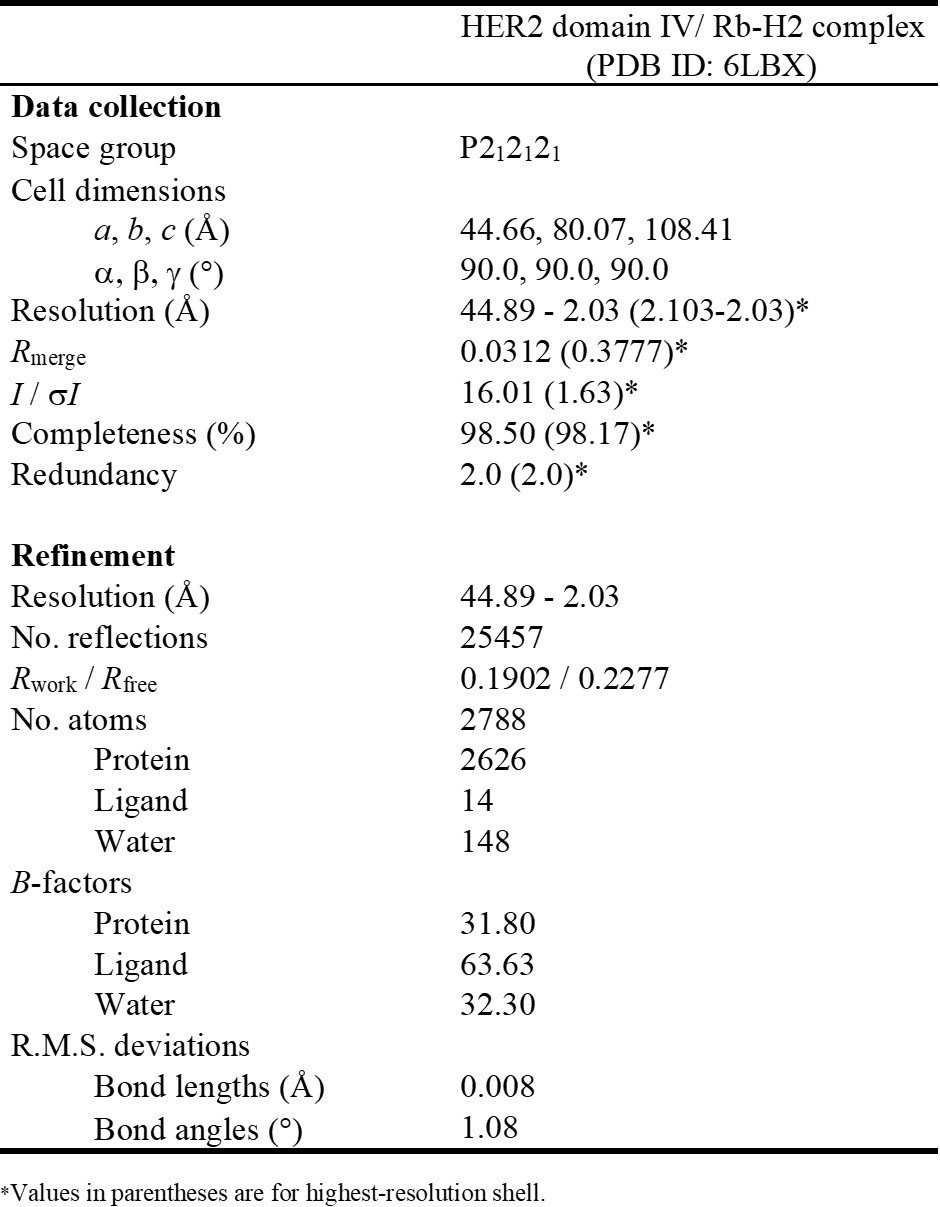
